# The Wild-type and Gain-of-function Mutant p53 Enhance p300 Autoacetylation through Conformational Switching

**DOI:** 10.1101/194704

**Authors:** Stephanie Kaypee, Raka Ghosh, Smitha Asoka Sahadevan, Shilpa Patil, Manidip Shasmal, Piya Ghosh, Neeladri Roy, Jayati Sengupta, Siddhartha Roy, Tapas K. Kundu

## Abstract

The transcriptional coactivator p300 is essential for p53 transactivation, although its precise mechanism remains unclear. We report that, p53 allosterically activates the acetyltransferase activity of p300 through the enhancement of p300 autoacetylation. Cryo-electron microscopy revealed that the domain organization of p300 is substantially altered upon binding of p53, suggesting that a structural switch may underpin the activation. Acetylated p300 accumulates near the transcription start sites accompanied by a similar enrichment of activating histone marks near those sites. Disruption of p53-p300 interaction by a site-directed peptide inhibitor abolished autoacetylated p300-mediated enhanced histone acetylation, suggesting a crucial role played by the allosteric activation in p53-mediated gene regulation. Gain-of-function mutant p53, known to impart aggressive proliferative properties in tumor cells, also activate p300 autoacetylation. The same peptide abolished many of the gain-of-functions of mutant p53 as well. We conclude that allosteric activation of p300 by p53 underpins gene regulation by p53. Reversal of gain-of-function properties of mutant p53 suggests that molecules targeting the p53-p300 interface may be good candidates for anti-tumor drugs.

## INTRODUCTION

Organismic complexity scales with the intricacies and sophistication of gene regulation. How this intricate regulation is achieved, however, is not fully understood. Specificity, both spatial and temporal, is at the heart of this extraordinarily elaborate and complex regulatory process (1). Protein-protein interactions among multiple transcription factors bound to adjacent sequences, DNA-mediated cooperativity and other factors have been found to enhance site selectivity, and to initiate the assembly of multi-protein regulatory complexes. After assembly the regulatory complex initiates the recruitment of coactivators, chromatin modifiers and other factors, thereby increasing chromatin accessibility and the initiation of RNA synthesis (2, 3). It is not known whether any steps beyond the assembly of regulatory complexes influence the spatial and temporal specificity of gene regulation.

p300 (KAT3B) is a transcriptional co-activator with intrinsic acetyltransferase activity. Being a large adaptor protein, p300 bridges the basal transcription machinery to DNA sequence-specific transcription factors (4, 5). By virtue of its structural plasticity, p300 can interact with the intrinsically disordered transactivation domains of a vast array of transcription factors, thereby functioning as a nodal integrator of signals terminating in regulation of transcription (6–9). Therefore, its recruitment via the interaction with sequence-specific transcription factors can regulate transcription from the chromatin template (10). This protein is often recruited to assembled regulatory complexes, where it acetylates histone tails in the vicinity. In this way, p300 promotes localized chromatin accessibility and, subsequently, selective transcriptional initiation (11).

The basal catalytic activity of p300 is enhanced by *trans*-autoacetylation of its lysine-rich regulatory loop (12). It had been proposed that in the absence of acetylation, the positively charged lysine-rich auto-inhibitory loop folds back onto the enzyme active site, thus hindering optimal substrate-enzyme interaction. Since p300 is involved in diverse physiological roles, the precise regulation of its function is essential. It has been demonstrated that some factors affect the levels of p300 autoacetylation in the cell. Intriguingly, these factors have distinct cellular functions and induce p300 autoacetylation under different cellular contexts. The Anaphase Promoting Complex/Cyclosomes (APC/C) subunits APC5 and APC7 are capable of enhancing p300 autoacetylation which is required for proper cell cycle progression (13). Similarly, the transcription factor, MAML1, has been shown to increase the levels of p300 autoacetylation thereby inducing the transcription of Notch pathway genes (14). During cellular stress, GAPDH associates with the E3 ligase, Siah1, and shuttles to the nucleus, where it enhances p300 autoacetylation which in turn activates the p53-mediated apoptotic pathway (15). p300 autoacetylation is negatively regulated by the Class III lysine deacetylase, SIRT2 and as expected, the suppression of p300 autoacetylation leads to the reduction of its enzymatic activity (16). We have previously reported the presence of global histone hyperacetylation in oral cancer. Mechanistically, this abnormality is a consequence of enhanced p300 autoacetylation, which is induced by NPM1 and nuclear GAPDH in a nitric oxide (NO) signaling-dependent manner. Ac-p300 may be the driving force that causes alterations in the epigenetic landscape and, consequently, the global deregulation of transcription that is required for oral carcinogenesis (17). Thus, modulation of p300 autoacetylation may be a key factor in regulating cellular responses.

p300 is a coactivator of the tumor suppressor p53 and acetylates it on 7 lysine residues, mainly on its C-terminal domain (18, 19). Studies have attributed the enhancement of DNA sequence-specific binding ability of p53 to the p300-mediated acetylation while contradictory findings have shown that p53 C-terminal acetylation may not have a profound effect on its promoter binding (20, 21). Nevertheless, accumulation of acetylated p53 has been observed during cellular stresses (22, 23) and p53 acetylation has been studied extensively with respect to cell cycle arrest, apoptosis, senescence and ferroptosis (24, 25). Studies using the acetylation-defective mutants of p53 have shown that the acetylation of p53 is not required for its DNA binding, rather it is crucial for the recruitment of co-activators such as p300 which are essential for p53 transactivation function (20). Thereby, reiterating the fact that the interaction between p300 and p53 is functionally important. p53 interacts with p300 in its tetrameric conformation, through its bipartite transactivation domain AD1 and AD2, while p300 interacts through several domains namely, TAZ1, KIX, TAZ2 and IBiD (26, 27). Intriguingly, numerous factors such as E1A, DDX24, WTX, MYBB1A, Skp2, can regulate p53 tumor suppressive functions *in vivo* by dictating its association with p300, thereby proving that the p300-p53 axis is an important cell fate determinant in stress responses (18, 28–32). The p53 gene acquires several mutations which abolish its tumor suppressive functions and confer oncogenic gain-of-function properties which contribute to tumorigenesis. Approximately 97% of the missense mutants map to the DBD of which a small number mutants occur in high frequency in cancers (33). Gain-of-Function (GOF) p53 mutants induce aberrant transactivation in cancers through the recruitment of p300, but the effect of mutant p53 on p300 catalytic activity is yet unknown. In the present study, we addressed the mechanism of factor-induced p300 autoacetylation and its importance in p53-dependent gene regulation.

We found that the master tumor suppressor protein p53 specifically induces p300 autoacetylation. To gain mechanistic insights into the induction of autoacetylation, we employed cryo-electron microscopy (cryo-EM) to generate 3D structures of free p300 protein as well as p53-p300 complex. Our cryo-EM data revealed that p300 undergoes substantial conformational rearrangements of its domains, thus leading it to adopt an active ‘open’ conformation following association with p53. Comparison of the genomic distribution of ac-p300 and p300 before and after stimulation by p53 indicates that ac-p300 is specifically and uniquely present around transcription start sites (TSS) of p53 regulated genes. This study establishes the mechanism of p53-mediated induction of p300 autoacetylation and its chromatin recruitment at transcriptional regulatory elements. Our data indicates that GOF mutant p53 may execute its downstream oncogenic functions through the modulation of p300/CBP autoacetylation and function. The implications of these observations are further explored.

## MATERIALS AND METHODS

### Antibodies

Primary antibodies used in this study: p300 (N-15), Santa Cruz (Catalog no. sc-584); p300 (C-20), Santa Cruz (Catalog no. sc-585); p53 (DO-1), Merck Millipore (Catalog no. OP43); Alpha-Tubulin (DM1A), Merck Millipore (Catalog no. 05-829). Rabbit polyclonal antibodies against autoacetylated p300 (K1499ac p300), H3K9ac, H3K14ac, H2AK5ac, H3 and GAPDH were raised in-house. Secondary antibodies: Goat Anti-Mouse IgG H&L (HRP), Abcam (Catalog no. ab97023); Goat Anti-Rabbit IgG H&L (HRP), Abcam (Catalog no. ab97051); Goat anti-Rabbit IgG (H+L) Cross-Adsorbed, Alexa Fluor^®^ 488, ThermoFisher (Catalog no. A-11008); Goat anti-Mouse IgG (H+L) Cross-Adsorbed, Alexa Fluor^®^ 633, ThermoFisher (Catalog no. A-21052).

### Cell Culture

Human non-small cell lung carcinoma H1299 cells (ATCC^®^ CRL-5803™) were cultured in RPMI-1640 media supplemented with 2 mM glutamine, 1% antibiotic solution (penicillin, streptomycin, amphotericin) and 10% fetal bovine serum (FBS). The pEBTetD p53 H1299 cells were cultured in supplemented RPMI-1640 under 1.2 μg/ml Puromycin antibiotic selection. Human hepatocellular carcinoma HepG2 cells (ATCC^®^ HB-8065™) and AW13516 cells, a human cell line derived from the Oral cavity squamous cell carcinoma (a kind gift from Dr. Amit Dutt (ACTREC, Mumbai, India)) were cultured in MEM supplemented with 2 mM glutamine, 1% antibiotic solution (penicillin, streptomycin, amphotericin) and 10% fetal bovine serum (FBS). The cells were grown at 37 °C in 5% CO_2_. Full length p300, PCAF and Tip60 were expressed in Sf21 (*Spodoptera frugiperda*) insect cells (ThermoFisher). The cells were grown in Graces’ Media containing antibiotic solution (penicillin, streptomycin, amphotericin) and 10% fetal bovine serum (FBS).

### Bacterial Cultures

The wild type p53 and p53 mutants (T18E p53, R273H, R175H, R249S, R248W, V143A, and L344A p53) were expressed and purified from BL21 (DE3) pLysS or BL21(DE3) *E. coli* strains.

### Cloning of p53 constructs

Wildtype p53 was cloned into BamHI-HF/HindIII-HF sites in pFLAG-CMV™-2 Mammalian Expression Vector (Sigma Aldrich). Point mutants were generated using Site Directed Mutagenesis (SDM) method using the QuikChange II XL Site-directed Mutagenesis Kit (Stratagene) according to the manufacturer’s protocol. The FLAG-tagged p53 construct was subcloned into the Tet-ON mammalian expression plasmid pEBTetD (a kind gift from Prof. Gründemann, University of Cologne, Germany (34)). The construct was amplified using a forward primer designed against the FLAG-tag containing a KpnI restriction site and a reverse primer specific to the 3’ end of p53 containing a XhoI restriction site. The primers have been listed in Table S3.

### Recombinant Protein expression and Purification

His_6_-tagged full length p300, Tip60 and FLAG-tagged PCAF were expressed by transfection of the respective recombinant baculovirus into Sf21 insect ovary cells. 60 hours post infection the cells were harvested. The His6-tagged proteins (His6-p300 and His6-Tip60) and FLAG-PCAF were purified as described previously (35, 36).

FLAG-p53 and FLAG-p53 mutants were expressed and purified from BL21(DE3)pLysS *E. coli* strain. The recombinant FLAG-tagged p53 proteins were purified through affinity chromatography using M2-agarose (Sigma Aldrich) binding resin as described previously (19).

### Small Molecule Inhibitor or Peptide treatment

HepG2 cells were treated with the small molecule inhibitor Nutlin-3a (5 μM) for 24 hours. The stabilization of p53 and status of p300 autoacetylation and acetylation of p300-substrates were determined by western blotting analysis and immunofluorescence. For the p53-p300 interfering peptides, HepG2 cells were treated with the indicated concentrations of peptides for 24 hours. The induction of wildtype p53 was achieved by treating the cells with Nutlin-3a for 24 hours after peptide treatment. Western blotting analysis was performed to study the effects of the peptides on p53-mediated modulation of p300 activity.

### RT-qPCR

Tet-inducible p53 expression H1299 cells were treated with doxycycline for 24 hours. The cells were harvested in Trizol reagent (Thermofisher) and RNA was isolated according to the manufacturers’ instructions. The purified RNA was treated with DNase I (New England Biolabs) at 37 °C for 20 min to remove any contaminating DNA. The RNA was again purified using Phenol-Chloroform-Isoamylalcohol (24:25:1) followed by ethanol precipitation at -80 °C overnight. cDNA was prepared using M-MLV Reverse Transcriptase (Sigma Aldrich) and oligo dT (23) primers (Sigma Aldrich). Real Time PCR was performed using SYBR Green (Sigma Aldrich) and mRNA-specific primers (Table S3).

### Co-immunoprecipitation

HepG2 cells were treated with 10 μM p53 phosphomimic peptide (3E peptide) or the scrambled control for 24 hours. The cells were then treated with 5 μM Nutlin-3a or the vehicle control (DMSO) for another 24 hours. The cells were harvested and lysed in RIPA buffer (150 mM NaCl, 50 mM Tris-Cl, 1% NP40, 1% sodium deoxycholate, 1 mM EDTA, and Protease inhibitors). p53 was immunoprecipitated using the DO1 antibody (Merck Millipore, Catalog no. OP43). Western Blotting analysis was performed to determine whether p53 and p300 interaction is disrupted by the p53 phosphomimic peptide.

### Wound Healing Assay

Cells were grown in a 30 mm dish or a 6-well plate. When the cells reach ~90% confluency, a scratch was created with a 10 μl tip held at a 45° angle. The cells were given a media change to remove all the floating cells. The wound created was monitored over a period of 24 hours to 48 hours till the wound closed. Images of the progress were taken at regular intervals.

### Immunofluorescence

H1299, HepG2, and AW13516 cells were grown on poly-lysine coated cover slips at 37 °C in a 5% CO_2_ incubator. The media was removed and the cell layer was washed in PBS to ensure all the media was removed. The cells were fixed in 4% paraformaldehyde for 10 minutes at room temperature. The cells were washed with PBS to remove the remnant PFA. The cells were then permeabilized in 5% Triton X-100. Cells were washed in PBS to remove the residual TritonX-100. The cells were blocked in blocking solution (5% FBS in PBS) at 37 °C for 45 minutes. The blocked cells were then incubated in primary antibody at the indicated dilution for 1 hour at room temperature on a reciprocal shaker. The cells were washed in washing buffer (1% FBS in PBS). The cells were then incubated in Alexa-fluor conjugated secondary antibody corresponding to the primary antibody used, incubated at room temperature for 1 hour. The nucleus was counter-stained with Hoechst 33258 (bis-benzamide (Sigma)), for 5 minutes at room temperature. Excess Hoechst stain was washed off with PBS. The cover slips were mounted in 70% glycerol onto a microscope glass slide. The stained cells were visualized using Carl Zeiss confocal microscopes LSM 510 META or the LSM 880 with Airyscan.

### Acetyltransferase Assays

For the p300, Tip60, and PCAF autoacetylation assay, the reactions were carried out in HAT assay buffer (50 mM Tris–HCl, pH 7.5, 1 mM PMSF, 0.1 mM EDTA, and 10% v/v glycerol) and 100 mM sodium butyrate at 30 °C for 30 min with or without the protein factors tested as autoacetylation inducers. The assay was performed in the presence of 1 μl 4.7 Ci/mmol [^3^H]-acetyl-CoA (NEN-PerkinElmer). After the autoacetyltransferase assay, the radiolabeled proteins were processed by fluorography. For the enzyme activity assay, reactions with 1 nM full-length p300 were carried out in HAT assay buffer (50 mM Tris–HCl, pH 7.5, 1 mM PMSF, 0.1 mM EDTA, and 10% v/v glycerol) and 100 mM sodium butyrate at 30 °C for 30 min. The assays were performed with 2 μM histone H3 in the presence of 1 μl 4.7 Ci/mmol [^3^H]-acetyl-CoA. After the HAT activity assay, the reaction mix was spotted on p81 phosphocellulose paper, which was washed in wash buffer (0.05 M sodium bicarbonate and 5 mM sodium carbonate). The spotted paper was dried and incubated in scintillation fluid (0.005% PPO and 5×10-4% POPOP in Toluene). The scintillation counts were measured in a scintillation counter (Wallac 1409 Liquid Scintillation Counter).

### Activity Rescue Assay

The 1 nM p300 was incubated in HAT buffer either at non-denaturating (30 °C, control) or at denaturating conditions (45 °C) for 15 minutes. The denatured enzyme was incubated in the presence of 80 nM and 200 nM p53 at 30 °C for 30 minutes. The substrate (2 μM recombinant histone H3) and [^3^H] acetyl-CoA were added to the reaction and the reaction was allowed to continue for 15 minutes at 30 °C. The reaction was stopped on ice followed by filter binding assay as described in the earlier section. The activity of p300 at non-denaturing conditions (30 °C) in the absence of NPM1 was considered as 100% activity.

### Cryo-Electron Microscopy

#### Sample for cryo-grid preparation

T18E p53, prepared in a buffer containing 10 mM Tris-Cl, 150 mM NaCl, 5 mM MgCl2, 20 μM ZnCl_2_, 2 mM PMSF, 1mM DTT, pH 7.5, was mixed with 200 nM p300 at a molar ratio 1:1 and incubated at room temperature for 5 min. For cryo-EM, 4 μl homogeneous p53-p300 complex was applied on glow-discharged Quantifoil^®^ holey carbon TEM grids (R2/2, Quantifoil, Micro Tools GmbH, Jena, Germany) coated with a home-made continuous thin layer of carbon followed by blotting and vitrification with a Vitrobot™ (FEI Inc, Hillsboro, Or, USA) and grids were frozen in liquid ethane. Purified p300, diluted to a final concentration of 200 nM in a buffer containing 10 mM Tris-Cl, 150 mM NaCl, 1 mM DTT, 2 mM PMSF pH 8.0, was used for sample preparation. Grids were prepared and sample imaged similarly as p53-p300 complex.

#### Cryo-EM data collection and 3D image processing

Data collection was performed on a Tecnai Polara microscope (FEI, USA) equipped with a FEG (Field Emission Gun) operating at 300 kV. Images were collected with 4K × 4K ‘Eagle’ charge-coupled device (CCD) camera (FEI, USA) at ~79000X magnification, resulting in a pixel size 1.89 Å at the specimen level with defocus values ranging from 1.5 to 4.5μm. All images were acquired using low-dose procedures with an estimated dose of ~20 electrons per Å^2^. Micrographs screening, particle picking were done separately with EMAN2 software (37) and SPIDER (38). Micrographs are selected on the basis of their visual quality, Thon rings in the power spectra and defocus values. Initially, for the complex, 5400 particle were picked manually according to their sizes using the BOXER program from the EMAN2 package with a pixel size of 1.89Å/pixel. Particle screening was done by visual inspection with utmost care so that only complex particles (~ 10-12 nm) were kept and particles of smaller sizes were rejected. 2D, 3D classification and initial model building were performed using EMAN2. The raw particle images were first subjected to multivariate statistical analysis and classification. The starting model was generated by the common line technique from selected 2D class averages. In SPIDER, defocus of each micrograph was estimated on the basis of its 2D power spectrum and then micrographs were divided into groups of similar defocus. Image processing was done using the reference based alignment method, where the initial volume of p53-p300 complex obtained from EMAN2 was used as the reference. Using projections of this model as a template, total 10,088 particles were selected semi-automatically from all the micrographs. The 3D reconstruction was then done following the standard SPIDER protocols for reference-based reconstruction (38). The overall resolution was 15.7 Å using the 0.5 cutoff criteria and 10.7Å using the 0.143 cutoff criteria. Data for the free protein were processed essentially as for the complex described above. Briefly, micrograph screening, particle selection, and 3D reconstruction were done in EMAN2 to generate an initial model. Coordinates of total 16,152 particles, picked manually in EMAN2, were imported in SPIDER for further reference-based particle alignment, 3D reconstruction, and refinement. The resolution of the final map was 13.5 Å using the 0.5 cutoff criteria and 9.8 Å using the 0.143 cutoff criteria.

#### Validation of initial models

Initial models were generated in EMAN2 from reference free 2D class averages. Validation of the 3D models was done in multiple ways. Reliability of the obtained initial model in EMAN2 was examined by comparing its different views with the reference-free 2D class averages produced in EMAN2, Xmipp (39), RELION (40), and reference-based SPIDER. Further, the 2D re-projections from the 3D maps were consistent with the 2D class averages. We also acquired two sets of images (grid preparation and data collection in different days) for both p300 and p300-p53 complex and processed independently resulting similar initial maps for both. The crystal structure of the Bromo-HAT shows close resemblance with the core structures of p300 in both the maps. We also manually produced another starting model from the crystal structure of the central domain (4BHW) by low pass filtering. 3D Image processing of the p300 dataset using two different starting models resulted similar 3D maps. Reprocessing of the image data of the complex using RELION (40), an alternative image processing software, resulted reconstruction fairly similar in overall structural features.

#### Model Building

There is no high resolution structure available for either p300 or p53 full length proteins. In order to describe the density maps of the p300 protein and p300-p53 complex in molecular terms we attempted to dock existing crystal structures of different domains of the protein. The detailed procedure is described in Supplementary Information section.

### Chromatin Immunoprecipitation

The cells were cross-linked in 1% formaldehyde. The crosslinking was quenched in 0.125 M glycine, the cells were lysed in SDS buffer, and the chromatin was sheared using a Bioruptor^®^ Diagenode instrument. Five to ten micrograms of antibody was used in each ChIP. The antibody and BSA-blocked protein G sepharose beads were added to the cleared sheared cross-linked chromatin and incubated at 4 °C overnight. The beads were then washed to remove non-specific binding with buffers containing different salt concentrations. The DNA-protein complexes were eluted from the washed beads using a SDS-sodium bicarbonate elution buffer. The eluates and input were decrosslinked at 65 °C for 6 hours. Proteinase K and RNase H were added to the decrosslinked lysates. The DNA was extracted using the phenol:chloroform:isoamyl alcohol method. The DNA was precipitated at -20 °C overnight using sodium acetate and glycogen in 100% ethanol. The DNA pellet was washed in 70% ethanol and dissolved in nuclease-free water. ChIP-qPCR was performed using SYBR Green (Sigma Aldrich) and loci-specific ChIP primers (Table S3).

### ChIP-seq and Bioinfomatic analysis

The Tet-ON p53 expression cell line was validated through Rt-qPCR for the expression of p53-responsive and non-responsive genes upon doxycycline treatment (Figure S5B). Libraries for ChIP-Seq were prepared with KAPA Hyper Prep Kits. The workflow consists of end repair to generate blunt ends, A-tailing, adaptor ligation, and PCR amplification. Different adaptors were used for multiplexing samples in one lane. Sequencing was performed on Illumina Hiseq3000/4000 for a single-end 150 run. High quality (HQ) reads with Q20 were generated using NGS QC Tool Kit v.2.3.3 (41). Filtered reads were aligned against *Homo sapiens* Ensembl build GrCh37 allowing 2 mismatches (-n 2) using the alignment tool Bowtie (42). Post alignment processing including SAM to BAM conversion along with sorting and PCR duplicate removal is performed using SAMTOOLS (43). BAM files of replicates were merged to get merged Input and IP BAM files for ac-p300 and p300. Peak calling is performed using MACS14 (44) using parameters mfold= 3:10 and p-value of 1e-5 (0.00001) (Figure S5C). The p300 and ac-p300 peaks were visualized using the Integrated Genome Viewer (IGV) tool (45). HOMER tool (46) was used for finding overlapping genes within distance of 100 bp overlap within ac-p300 and p300 and to find unique genes specific to ac-p300 and p300. Total number of peaks obtained were annotated using Peak Analyser v1.4 (47). Bedgraph files from the peak bed file generated using BEDTOOLS sortBed and GenomeCoverageBed commands. Integration of gene expression data from microarray analysis with ChIP-seq gene enrichment data is carried out to check for the binding occupancy and gene regulation. Average profiles and heatmaps were generated showing ChIP signal densities spanning 5 kb upstream and downstream of TSS using ngs.plot.r (48).

### Statistics

Unpaired two-tailed Student t-test has been used for the statistical analysis. The p-values have been indicated in the figures, legends, and methods.

## RESULTS

### p53 directly modulates p300 autoacetylation and acetyltransferase activity

To explore whether p53 has an effect of acetyltransferase activity of p300, an *in vitro* acetyltransferase assay was designed to detect the effect of p53 on the levels of p300 autoacetylation. p53 specifically induced the autoacetylation of 20 nM p300 in a concentration-dependent manner (ranging from 40 nM to 160 nM p53 monomers) but exhibited a negligible effect on the autoacetylation of the other two classes of KATs, PCAF and Tip60 (Figure 1A). An approximately 4.5-fold enhancement of p300 autoacetylation was observed in the presence of 160 nM p53 (4:1 ratio of p53 tetramers to p300). Because p53 forms a stable tetramer and may possibly interact with p300 as a tetramer (27), a significant degree of p300 autoacetylation was reached only when p53 was present in a 1:1 ratio of tetrameric p53 to p300. At a suboptimal concentration (40 nM) of p53, (tetrameric p53 to p300 ratio of 0.5:1), a negligible increase in the autoacetylation was observed (Supplementary Figure S1A). Moreover p53 is a substrate of most major nuclear KATs, including p300/CBP (19), PCAF (KAT2B) (49), TIP60 (KAT5) (50, 51), MOZ (KAT6A) (52) and MOF (KAT8) (50). Therefore, the specificity exhibited by p53 toward the enhancement of p300 autoacetylation, with no effect seen for PCAF and Tip60, led us to conclude that p53 is not merely a substrate of p300 but a regulator of the intrinsic enzymatic activity of p300. This specific role of p53 was investigated further. Three different substrates of p300, p47 (N-terminal truncated (Δ40p53) isoform of p53, incapable of binding to p300), Positive Coactivator 4 (PC4) and p50 (a subunit of NFκB) were also tested and found not to effectively induce p300 autoacetylation (Figures 1B-1D). To investigate the effect of the p53-mediated induction of p300 autoacetylation, we performed an *in vitro* histone acetyltransferase assay using recombinant histones as the substrate. In agreement with the results of the autoacetylation assay, we observed an approximately 3-fold increase in p300 histone acetyltransferase at a p300: p53 in a molar ratio of 1:2. Notably, we observed an approximately 4.5-fold increase in p300 activity in the presence of p53, thus suggesting that p53 positively modulates p300 KAT activity through the induction of autoacetylation. Further increases in the p53 concentration did not affect the HAT activity beyond 4.5-fold (Figure 1E). The *in vitro* p53-mediated induction of p300 autoacetylation was further verified in the cellular context. Endogenous p53 levels were stabilized in HepG2 cells by using the small molecule inhibitor of the MDM2-p53 interaction, Nutlin-3a. The stabilization of p53 after Nutlin-3a treatment appeared to positively regulate the levels of p300 autoacetylation in the cells in comparison to DMSO-treated or untreated cells (Figure 2A). Furthermore, the fluorescence intensity due to p300 autoacetylation was significantly (p<0.001) enhanced in HepG2 cells treated with Nutlin-3a expressing wild-type p53 (Figure 2B and 2C). We expected that hyper-autoacetylated p300 would induce p300-specific histone acetylation in cells. To address this possibility, histone acetylation at H3K9 and H3K14 were investigated in the presence and absence of Nutlin-3a (Figure 2D). It has been previously demonstrated that p53 acetylation and histone acetylation increase after Nutlin-3a treatment (53). Therefore, to test for whether the enhancement in histone acetylation was due to the p53-mediated enhancement of p300 autoacetylation, we treated p53-null H1299 cells with Nutlin-3a and probed for the levels of different histone acetylation marks in treated versus control cells (Figure 2E). Interestingly, we found that the histone acetylation levels did not show any appreciable alteration in Nutlin-3a-treated H1299 cells, thereby signifying the importance of p53 in the activation of p300 KAT activity. To rule out the possibility that p300 might be transcriptionally regulated by p53 rather than post-translationally modulated, p300 transcript levels were compared between cells treated with Nutlin-3a and DMSO. No change in *EP300* transcript levels was observed in the presence of p53 compared to *CDKN1A* (an early p53 target gene) transcripts, thus suggesting that p300 is not regulated by p53 at the transcriptional level (Figure 2F). Collectively, these results established p53 as a *bona fide* inducer of p300 autoacetylation and its acetyltransferase activity. To elucidate the mechanistic details of p53-mediated activation of p300, the importance of the direct interaction between p53 and p300 was investigated. p53 directly contacts p300 through four important residues (L22, W23, W53, F54) in its transactivation domain. Mutations in these residues severely compromise p53 transactivation (27). After DNA damage, p53 is phosphorylated by several kinases, and phosphorylated p53 binds to its coactivator p300 with higher affinity (54, 55). Interestingly, the higher the number of phosphorylated sites on the p53 N-terminus, the higher its affinity for CBP/p300 (55, 56). A triple-phospho-mimic peptide (S15E, T18E, S20E) comprising of the 11^th^ to 30^th^ amino acid residue of the p53 (TAD1 domain) protein along with a cell penetrating sequence and NLS was used to obstruct the interaction between p53 and p300 (57) (Supplementary Figure S1B, Table S1). Using a filter-binding assay-based strategy, the importance of the p53-p300 interaction for the activation of p300 was assessed on the basis of the level of p300-mediated histone acetylation (Figure 2G). The peptides and p53 alone did not bind to the p81 phosphocellulose filters; therefore, the measured radioactivity was only from the tritium-labeled acetylated histones bound to the filters. The basal activity of p300 in the absence of any inducer was considered as 100% activity, and the peptides alone did not alter the activity of p300, thus signifying that any change in histone acetylation observed thereafter was a direct consequence of the p300-p53 interaction. In the presence of 4 nM p53, a 3-fold increase in the histone acetylation was observed. The presence of scrambled peptide did not alter the p53-mediated induction of p300 activity, but triple phospho-mimic p53 peptides (3E pep) effectively decreased histone acetylation levels. These results indicated that the 3E phospho-mimic p53 peptide effectively interfered with the p53-p300 interaction, thus presumably leading to decreased p300 autoacetylation and activity, as indicated by a significant decrease in histone acetylation levels. The scrambled peptide exhibited a negligible effect on p53-mediated p300 activation, thus ensuring that the decrease observed in the case of the phospho-mimic peptide was indeed due to the disruption of the p53-p300 axis and was not a non-specific effect of the peptides (Figure 2G). Furthermore, to ascertain whether these peptides were effective in cells, HepG2 cells were treated with Nutlin-3a to stabilize the levels of p53. The cells were then treated with different concentrations of the 3E phospho-mimic p53 peptide, which has cell-penetrating sequences (Figure 2H). The levels of p53 increased after Nutlin-3a treatment, and the peptide treatment did not alter p53 protein levels. Therefore, the mimic p53 peptides did not alter cellular p53 protein levels, and the observed alteration in histone acetylation is a direct consequence of the disruption of p53-p300 interaction. In agreement with the previous observation, the H2AK5 acetylation increased on p53 stabilization with Nutlin-3a (Figure 2H and Supplementary Figure S1C). After treatment with the interfering peptide, the levels of H2AK5 acetylation gradually decreased in a concentration-dependent manner (Figure 2G), whereas H2AK5ac levels remained unchanged after scrambled peptide treatment (Supplementary Figure S1C). The direct interaction between p300 and p53 appeared to be crucial for the induction of p300 catalytic activity; therefore we wanted to investigate whether p53 could modulate the structure of p300. Since p300 is a large, intrinsically disordered protein of 300 kDa, it could be a possibility that p53 may act as a molecular chaperone for p300, thereby stabilizing its disordered structure. To address this, we designed an in vitro radioactivity-based histone acetyltransferase assay where we denatured p300 at 45 °C and then incubated the denatured enzyme with p53 at 30 °C. The refolding or recovery of p300 activity was scored by its ability to acetylate recombinant histones. Interestingly, we obverse a p53 concentration-dependent rescue of p300 activity, suggesting that p53 could possibly act as a molecular chaperone for p300 (Figure 2I). Overall, we observed that p53 is a potent inducer of p300 autoacetylation and enzymatic activity which may be through the modulation of p300 structure by p53.

**Figure 1:**
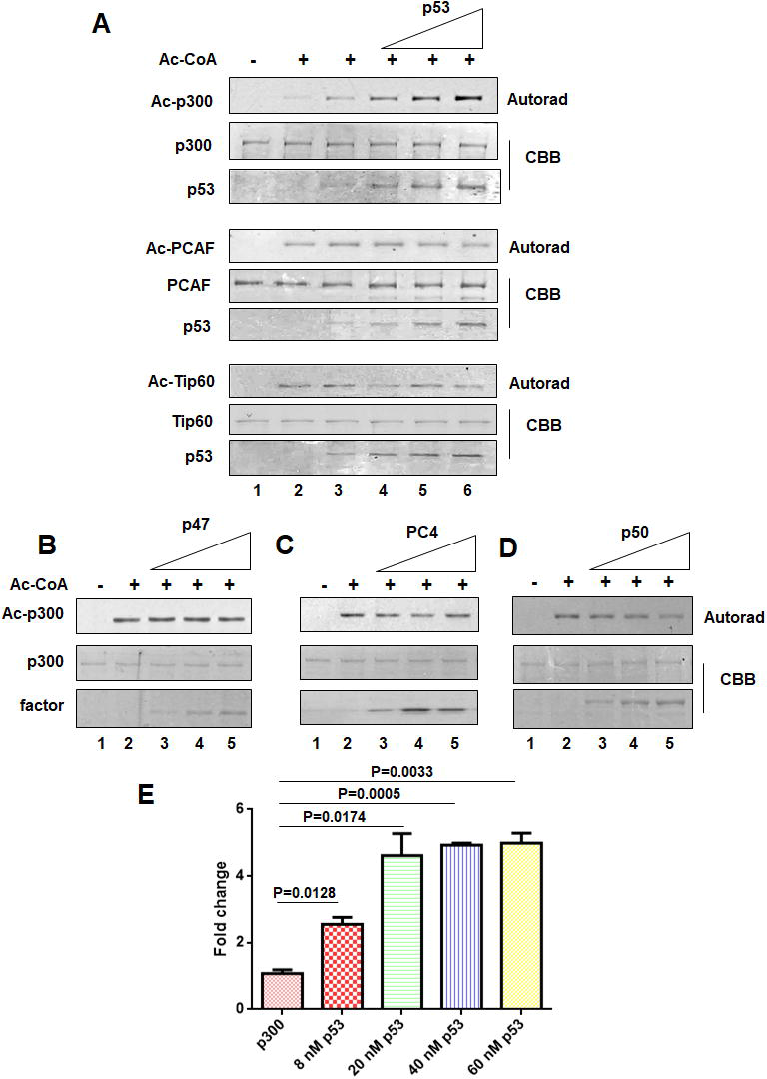
The tumor suppressor p53 enhances p300 autoacetylation. (A) An *in vitro* acetyltransferase assay was performed to determine the levels of autoacetylation of different lysine acetyltransferases (KAT) in the presence of recombinant p53. The autoradiograms indicate the levels of p300, PCAF, and Tip60 autoacetylation respectively; while the Coomassie stained protein panels show the loading control. (B-D) An *in vitro* acetyltransferase assay was performed to determine the levels of p300 autoacetylation in the presence of different p300 substrates (B) p47, (C) PC4 and (D) p50. (E) Enhancement of p300 catalytic activity in the presence of increasing concentrations of p53 as indicated was determined by filter binding histone acetyltransferase assay, using recombinant histone H3 as substrate. The mean fold change was plotted ± SD, statistical analysis was performed using two-tailed unpaired Student t-test.

**Figure 2:**
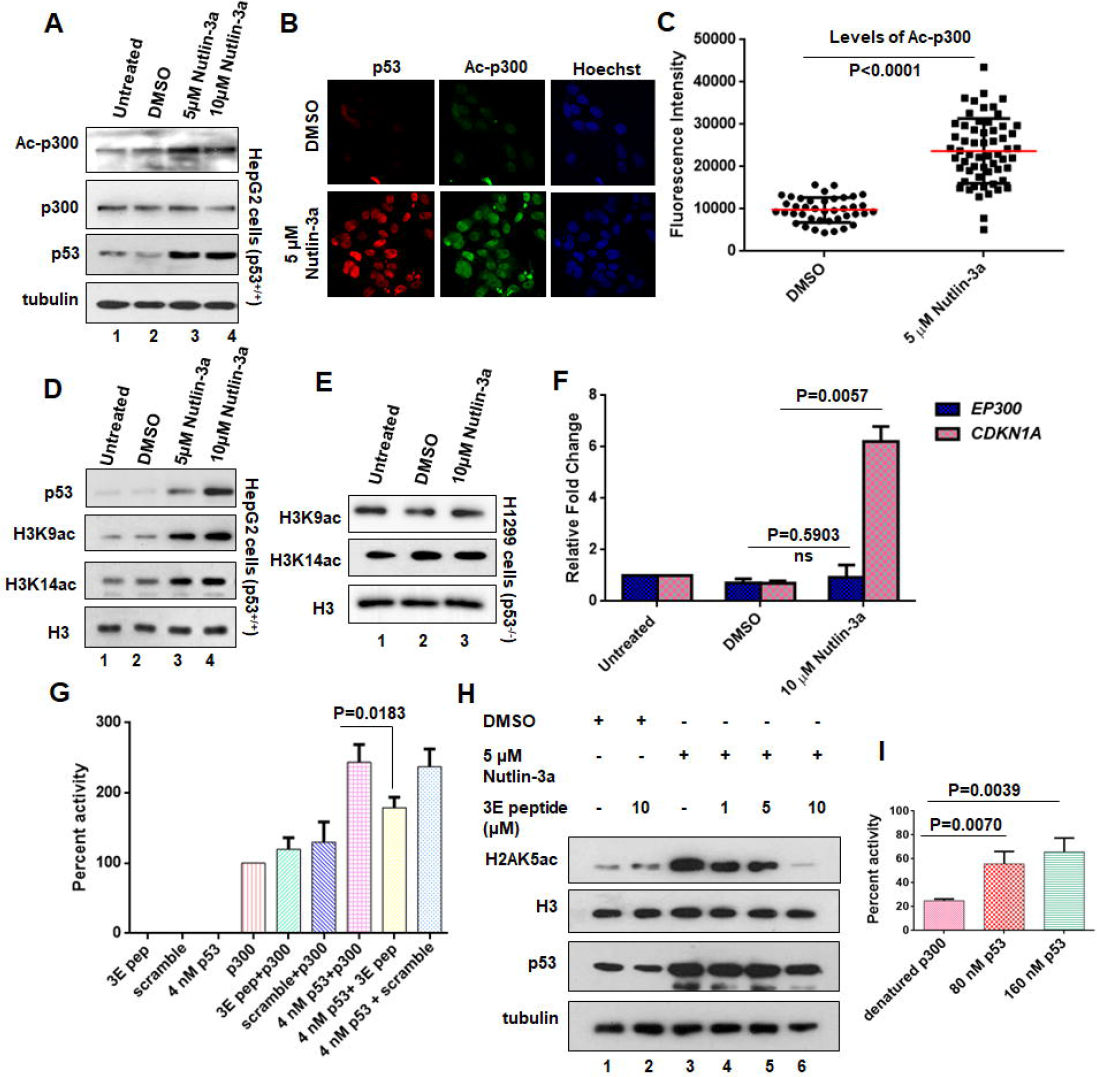
p53 modulates p300 autoacetylation and acetyltransferase activity in cells. (A) HepG2 cells were treated with DMSO (vehicle control) or Nutlin-3a as indicated (Lane 2-4), levels of ac-p300, p300, p53 were analyzed by immunoblotting. Alpha-tubulin levels were considered as the loading control. (B) HepG2 cells were treated with 5 μM Nutlin-3a for 24 hours and the levels of autoacetylated p300 (ac-p300) and p53 were assessed by co-immunofluorescence. (C) The fluorescence intensity of ac-p300 in control cells versus Nutlin-3a treated cells have been quantified (represented as mean ± SD, unpaired two-tailed Student t-test, n=50 of three independent experiments). (D and E) HepG2 cells (D), and H1299 cells (E), were treated with DMSO (vehicle control) or Nutlin-3a as indicated. Histone acetylation levels were analyzed by immunoblotting using H3K9ac and H3K14ac antibodies. Total histone H3 levels were used as the loading control. (F) The relative transcript levels of *EP300* and *CDKN1A* (p53-responsive gene) were determined by qRT-PCR (mean ± SD). Unpaired two-tailed Student t-test statistical analysis was performed. Actin transcript level was used as the internal control. The relative p300 transcripts do not alter significantly on Nutlin-3a treatment. (G) Results of filter binding assay indicate that the phosphomimic p53 N-terminal peptide can effectively reduce the activity of p300 by interfering with its interaction to p53. (H) HepG2 cells treated with Nutlin-3a or DMSO, and p53 phosphomimic peptide (3E pep) as indicated, and the levels of H2AK5ac, H3, p53, and tubulin were probed by immunoblotting. (I) The rescue of heat-denatured p300 activity (activity rescue assay), upon addition of wild type p53, at the indicated concentrations, was determined by the *in vitro* filter-binding assay, using recombinant histone H3 and [^3^H] acetyl-CoA.

### Cryo-EM reveals conformational changes in p300 following p53 binding

To elucidate the mechanism of p53-mediated structural alteration and activation of p300, we employed cryo-electron microscopy (cryo-EM) to visualize the full-length p300-p53 structure. A plausible hypothesis is that p53 binding stimulates p300 activity through a structural switch, resulting in an active p300 conformation. Therefore, to gain mechanistic insights into the induction of autoacetylation, we have generated the 3D structures of free p300 protein as well as p53-p300 complex. We have generated cryo-EM maps depicting p300 alone (Figure 3A and 3B) and in complex with p53 (Figure 3C and 3D) (obtained at resolutions of 9.8 Å and 10.7 Å at FSC = 0.143 cut off, and 13.5 and 15.7 Å at FSC = 0.5 cut off for p300 and the p53-p300 complex respectively; Supplementary Figure S2R and S2Q). To ensure the reliability of the maps, we used several validation strategies (see Materials and Methods; Supplementary Figure S2A-S2P).

**Figure 3:**
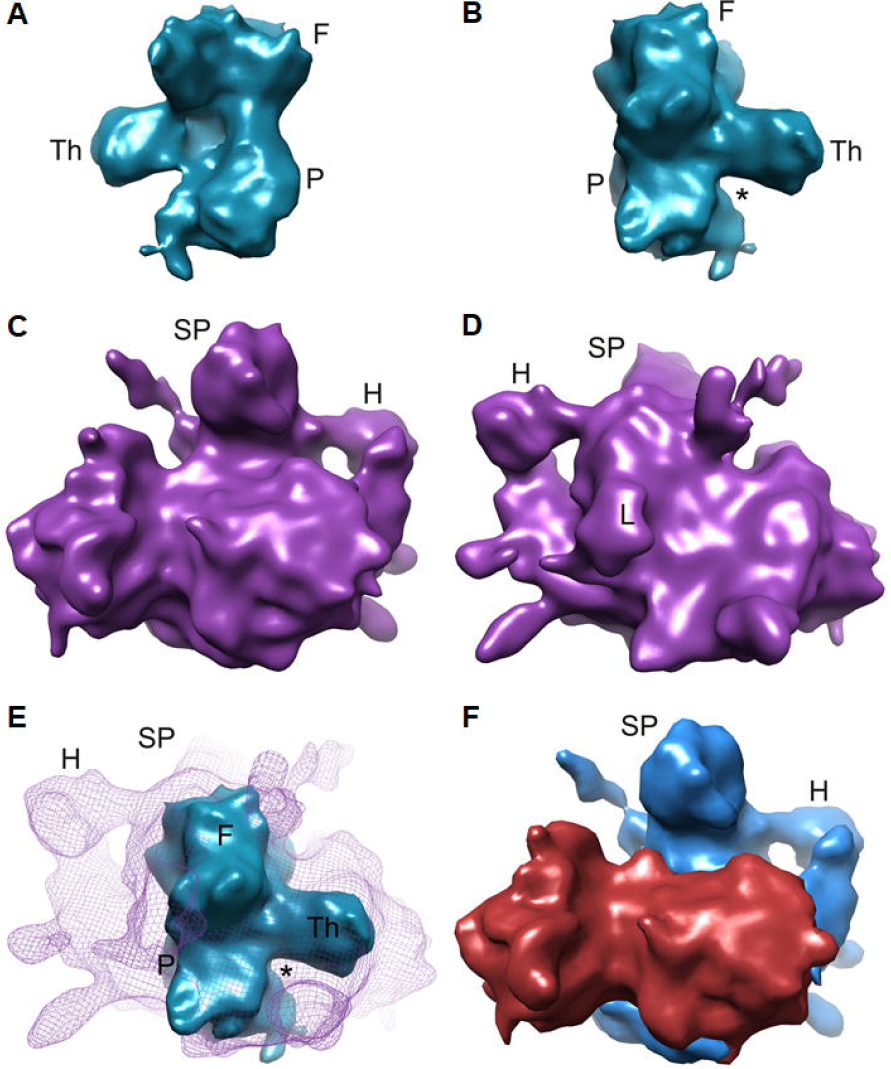
Cryo-EM 3D reconstructions of p300 and p300-p53 complex. (A) & (B), Front (A), and back (B) view of 3D cryo-EM map of p300 (surface representation in dark cyan). (C) & (D), Front (C) and back (D) view of the density map of p300-p53 complex (purple surface). Land marks in p300: F, fist; Th, thumb; and P, palm. In complex: SP, spout; H, handle; L, loop-like structure, density attributed to autoinhibitory loop. (E) Superimposition of the p300 map (dark cyan) on the p300-p53 complex map (purple mesh) reveals that substantial conformational changes occur in p300 upon p53 binding. The asterisk (*) indicates absence of density in the p300 map, whereas in complex, density can be clearly seen in this region. (F) Surface representation of the densities encompassing p300 (blue) and p53 (brick red) in the complex following segmentation.

On the basis of the size of the enclosed volume, the molecular mass of the p300 density map (~233 kDa estimated from the occupied volume, with 0.82 D/Å^3^ as the protein density) appeared somewhat smaller than the known molecular mass of p300 (264 kDa). Evidently, the long unstructured regions at the N- and C-termini (~650 amino acids, equivalent to ~70 kDa; Supplementary Figure S2T) were not entirely visible in our reconstruction, likely because of their dynamic nature. Considering that the p300 map encompassed only the structured domains, the effective molecular mass was adequate. No high resolution structure of the full-length p300 is available so far. Only a cryo-EM study of the estrogen-coactivator complex has recently reported (58) that p300 forms a rather compact yet flexible conformation when present outside the complex, thus indicating the intrinsically disordered nature of the protein. Although the resolution of the published map is much lower compared to the map we present here, some of the features of our p300 reconstruction could be matched with the previous map (58). However, the size of the previous map is much larger, thereby suggesting that the density of the previous map presumably encapsulates almost the entire protein.

The p300 density map showed a ‘fist-like’ appearance with a ‘thumb’ feature (Figure 3A and 3B), whereas the p53-p300 complex adopted a shape similar to that of a kettle, with a protruded ‘spout’ in the middle and a ‘handle’ at the side (Figure 3C and 3D). Clear resemblance of the overall topology of the free p300 density with part of the p53-p300 complex density map could be identified when the two maps were juxtaposed (Figure 3E). However, compared with the p300 in complex with p53, the free protein appeared more compact. A bi-lobed density attributable to the p53 tetramer was visible within the density map of the complex (Figure 3F). The segments corresponding to p300 and p53 densities were computationally separated by using a cluster segmentation procedure implemented in SPIDER (38). Comparison of the two maps indicated conformational alteration in p300 domains following p53 binding. The overall topology of the p53-p300 complex (Figure 3F) appeared to be very similar to that of the predicted model in a previous structural study (27), the centrally located, protruding density (termed here as ‘spout’) being the structural hallmark in the density map of p300 in complex with p53 (Figure 3C, 3F and Supplementary Figure S2S).

### Structural modulation and activation of p300 upon allosteric binding of p53

A prediction regarding the elusive ‘active’ p300 conformation has been made in a previous analysis of the p300 core domain crystal structure by Delvecchio *et al*, in which the authors have suggested that free p300 may be present in an inactive conformation (59). However, the molecular events underlying factor-induced p300 autoacetylation are not clearly understood. To dissect the mechanistic details of the p53-induced activation of p300 autoacetylation, we attempted to interpret the cryo-EM densities in molecular terms. Available atomic structures of different domains were used to identify the location of this domain within the EM density envelopes. The bromo-HAT-RING-PHD domains of p300 comprise the central domain, while the KIX and TAZ1 domains are at the N-terminal, and the TAZ2 and IBiD domains belong to the C-terminal part (Supplementary Figure S2T). The largest piece of the p300 protein that could be crystallized is the central domain which shows an ‘auto-inhibited’ conformation (PDB: 4BHW), where PHD and RING domains are loosely packed on the HAT domain. The structure reveals a ‘kink’ between the bromo and HAT domains (Supplementary Figure S2V). Although the resolution of the maps are limited, in agreement with the previous prediction (59) our structural data clearly manifested that the central region of p300 density (Fig. 3E) transforms to an ‘open’ conformation following association with p53.

Remarkable resemblance of the central domain crystal structure with the free p300 density map was identified. The crystal structure (4BHW) when fitted in Chimera (60) as a rigid body into the free p300 density map (CC = 0.88), it could satisfactorily be accommodated in the central region of the density (Figure 4A and Supplementary Figure S2W). The HAT occupied the ‘palm’ while bromo domain was placed in the ‘fingers’ of the ‘fist’ (Supplementary Figure S2V and S2W) keeping the ‘kink’ between the HAT and bromo domain unaltered.

**Figure 4:**
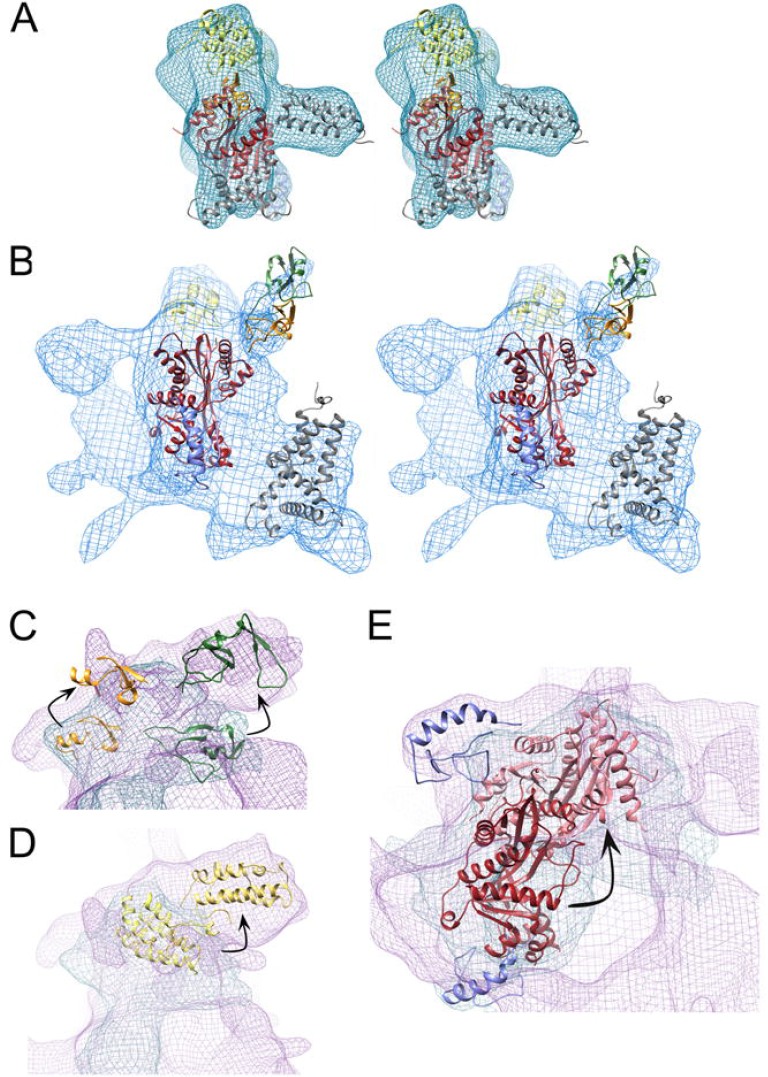
Interpretation of the densities in molecular terms. (A) & (B), Stereo representation of (A) p300 map (cyan mesh) and (B) the difference map corresponding to the p300 density obtained from the complex map (blue mesh). Both maps are fitted with crystal structure of various domains of p300 (PDB: 5HOU, 4BHW, 2LQH, and model of autoinhibitory loop generated by PHYRE2). Bromo, RING, PHD, HAT domains, and autoinhibitory loop are shown in yellow, green, orange, red and violet respectively and other p300 domains are colored grey. (C-E) Close up views of the superposed cryo-EM maps of p300 and p300-p53 complex (blue mesh: p300 map; purple mesh: p300-p53 complex map) showing rearrangement of (C) RING (green) and PHD domain (orange); Conformational changes in (D) Bromo (yellow) and (E) HAT domain (red) along with autoinhibitory loop (violet) following complex formation with p53.

On the other hand, the protruded feature (‘spout’) in the density map of p300 in complex with p53 (Figure 3C and Supplementary Figure S2S) showed striking resemblance with the shape of the bromo domain (Supplementary Figure S2S and S2V). However, to accommodate the bromo domain into that density, the opening of the ‘kink’ between HAT and bromo domain was obligatory (Figure 4B, Supplementary Figure S2S). Thus, the p300 in the p53-p300 complex manifested its ‘open’ conformation revealing the protruded shape (‘spout’) of the bromo domain prominently (Figure 4D). Furthermore, the densities corresponding to the RING and PHD domains occupied the density at the junction of ‘fist’ and ‘thumb’ in the p300 map (Supplementary Figure S2U-S2W), whereas no such densities were visible in the same region of the complex (Supplementary Figure S2S). Instead, weak densities were observed at the top of the ‘spout’ (Figure 4B and Supplementary Figure S2S). Apparently, the PHD and RING domains were displaced (Figure 4C) in a similar manner suggested in a recent study (59).

On the basis of the cylindrical shape of KIX (the N-terminal domain preceding the bromo domain), the ‘thumb’ density was assigned to KIX (Supplementary Figure S3A). Whereas, a triangular shaped density resembling the structure of the TAZ domain apparently fastened the HAT domain in a ‘closed conformation’ of the protein which opened up in the complex (Supplementary Figure S3B). Overall, it appeared that, in addition to the RING domain, the ‘auto-inhibited’ inactive conformation described in the crystal structure (Delvecchio et al., 2013) was further stabilized by other domains of the protein. A clear loop-like density was seen at the back of the HAT domain in the complex (Figure 3D and 4E). A model of the lysine-rich auto-inhibitory loop could be placed inside this density after a rotation along the HAT domain axis. We speculate that, while transforming into the ‘open’ conformation upon interactions with p53, the HAT domain of p300 rotates and exposes the auto-inhibitory loop (Figure 4E).

Distinct features of the p53 tetramer were identifiable in the cryo-EM map of the p53-p300 complex. In the previously analyzed p53 tetramer structures (without DNA or DNA-bound maps, approximately 6-8 nm, the four DNA-binding domains remain closely packed (61, 62), while the tetramerization domains are displaced out of the plane (Figure S4). In contrast, in the present map of p53-p300 complex (approximately 12 nm), the p53 tetramer adopted a distorted shape (Supplementary Figure S3C), with two side-lobes exhibiting a centrally connected density attributable to the packed tetramerization domains (PDB: 4D1M; Supplementary Figure S3E). A dimer of p53 DNA-binding domains (presumably along with the N-terminal domains) could be accommodated into each side lobe of p53 density (Supplementary Fig S3C). A segment of DNA could be accommodated in the cleft created by the two side lobes (Supplementary Figure S3D).

Overall structural arrangements of the p53-p300 complex demonstrated striking similarity to a previous predicted model where it has been proposed that the p300 protein wraps around the p53 tetramer. In this model, the N-terminal KIX and TAZ1 domains and the C-terminal TAZ2 and IBiD domains of p300 interact with p53, with the TAZ2 interaction being the strongest, followed by the interactions with TAZ1, KIX, and IBiD (27). Thus, our map of the p53-p300 complex (CC with the density envelope = 0.9) not only provided experimental evidence of this proposed model but also elucidated the dynamic mechanism of factor-induced p300 autoacetylation.

### Redistribution of autoacetylated p300 upon p53-mediated activation

If the p53-mediated allosteric activation of p300 is important for p53 mediated regulation of targeted genes, then it is expected that activated forms should be present in the vicinity of the regulated genes. We have thus attempted to explore the effect of p53 expression on presence p300 in different genomic sites. The differential occupancy of ac-p300 and p300 on the promoters of genes that are expressed after p53 stimulation has been investigated (Supplementary Figure S5C). The enrichment patterns indicated a distinct preference of ac-p300 for the promoter proximal regions, after p53 expression, whereas only a weak enrichment pattern was observed for p300 (Figure 5A and Supplementary Figure S5A), thus reinforcing the possibility that the autoacetylated form of p300 is actively associated with p53-mediated transcriptional initiation. We analyzed the differential occupancy of p300 and ac-p300 with publically available datasets of RNAPII, H3K27ac and H3K4me1 distribution. The abundance of ac-p300 at the promoters correlated with the presence of RNA Pol II enrichment and active promoter marks such as H3K27ac (Figure 5A). These data indicated a specific role for ac-p300 within the entire pool cellular p300. The coactivator occupancy has been integrated with the microarray expression analysis of the p53-induced differentially expressed genes (25) (Supplementary Figure S5B), revealing a greater overlap between the p53-driven transcription and ac-p300 (4341 genes) recruitment (Figures 5C and 5D) in comparison to p300 (1950 genes) (Figure 5B).

**Figure 5:**
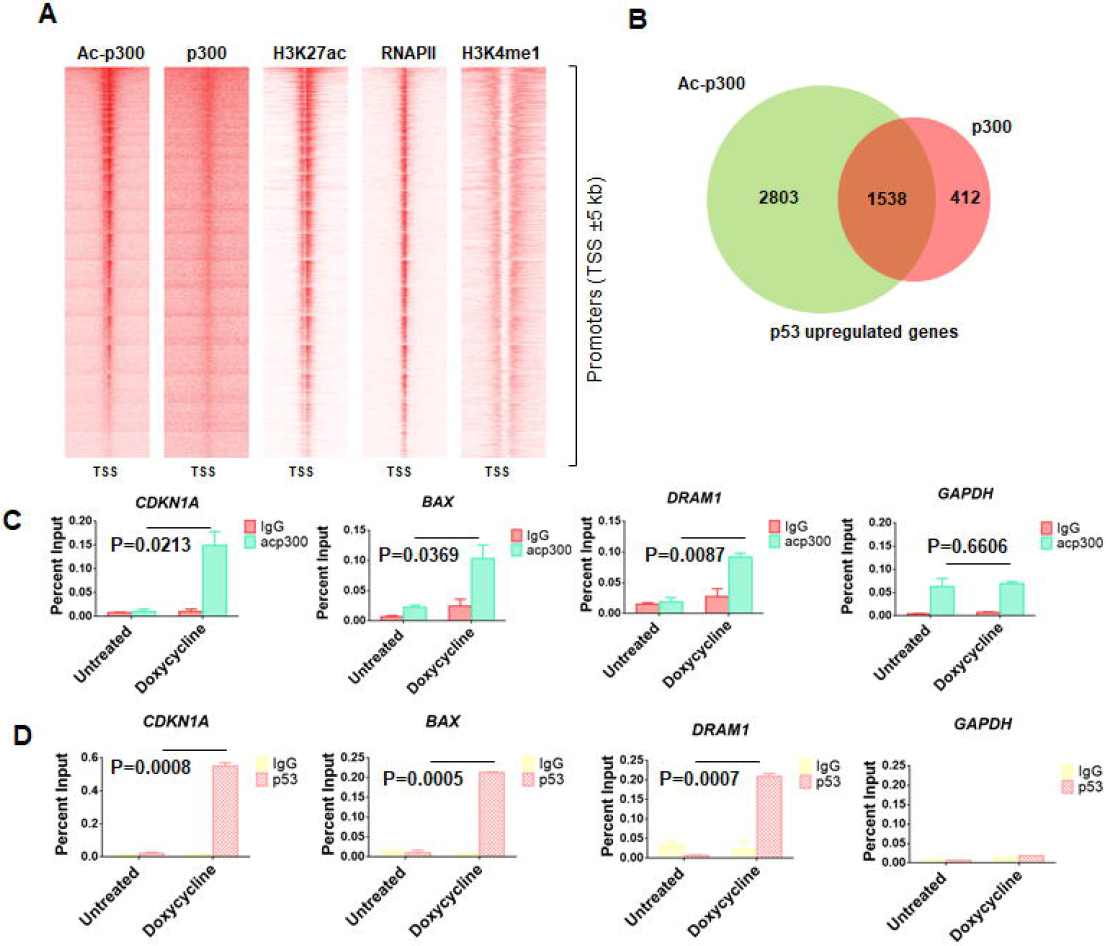
Redistribution of p300 after p53-mediated enhancement of p300 autoacetylation. (A) Heatmaps depicting the enrichment of ac-p300, p300, H3K27ac, RNA Polymerase II (RNAPII), and H3K4me1 at promoters (TSS±5 kb). (B) Venn diagram showing the overlap between the up-regulated genes enriched at the promoters with both ac-p300 and p300 and genes unique to each set. (C) & (D), ChIP-qPCR to determine the occupancy of (C) p53 and (D) autoacetylated p300 (ac-p300) at p53-responsive gene promoters.

### The Gain-of-Function (GOF) mutant p53 induces p300 autoacetylation to enhance tumorigenesis

The *TP53* gene is mutated in approximately 50% of malignancies, of which 75% are missense mutations (33, 63). The missense mutations mapping to the DNA binding domain not only abrogate the tumor suppressive functions of p53 but also impart unique oncogenic functions to the mutant protein (33). Since our previous results elucidated the mechanisms of p53-mediated induction of intermolecular p300 autoacetylation and the role of autoacetylated p300 in p53 downstream gene expression, we investigated whether these GOF mutants retained the ability to enhance p300 catalytic activity through the induction of autoacetylation. We tested different GOF hotspot mutants, three conformational mutants (R175H, V143A, and R249S) and two DNA contact mutants (R273H and R248W), in an in vitro autoacetylation assay. All the GOF mutants appeared to enhance p300 autoacetylation in vitro which was comparable to the enhancement observed in the presence of the wild type p53 (Figure 6A). In H1299 cells ectopically expressing GOF p53 mutants, the levels of autoacetylated p300 were checked by immunofluorescence in the mutant p53 transfected cell versus the untransfected cells. The intensity of autoacetylated p300 immunofluorescence staining revealed that the cells expressing the GOF mutants, both DNA contact mutants (Supplementary Figures S6A and S6B) and conformational mutants (Supplementary Figure S6C and S6D), appeared to have significantly higher levels of autoacetylated p300 in comparison to the untransfected controls (Supplementary Figure S6). To understand the pathophysiological relevance of GOF mutant p53-mediated enhancement of p300 autoacetylation, the R273H p53 mutant, which has one of the highest incidences in cancers (6.7%), was chosen for further investigation. To establish that R273H p53 mutant could alter the levels of p300 autoacetylation alone without affecting the overall p300 protein levels, immunofluorescence was performed in H1299 cells ectopically expressing R273H p53. As observed in the earlier immunofluorescence staining, the autoacetylation of p300 increased dramatically in the presence of R273H (Figure 6B, upper panel) while exhibiting a negligible change in the overall p300 protein levels (Figure 6B, lower panel). These data reinforce the fact that similar to wild type p53, the GOF R273H mutant too, modulates p300 only at the autoacetylation level and does not appear to affect p300 protein stability or expression. As established earlier, that increased KAT activity is a direct consequence of p300 autoacetylation, the levels of different p300-specific histone marks were checked, after ectopically expressing the mutant R273H and wild type p53 as a control. It is evident from the western blotting analysis, that histone acetylation on H2AK5 and H3K9 were upregulated in cells transfected with the mutant and wild type p53 in comparison to the vector-transfected cells (Figure 6C). To investigate the role of R273H mutant-mediated induction of p300 autoacetylation had a role in regulating the tumorigenic potential of the mutant, an inducible Tet-ON H1299 cell line for R273H p53 expression was created. Earlier experiments have established that the direct interaction of p53 with p300 is essential for the induction of p300 autoacetylation. Therefore, the aim of the experiment was also to test whether the direct interaction of p300 interaction the p53 mutant R273H played a role in the GOF of this mutant. In a wound healing assay, the cells were treated with doxycycline to induce the expression of R273H p53 and the untreated cells served as the experimental controls. Under both conditions, R273H p53 presence and absence, the cells were treated with either the p53-p300 interfering phosphomimic p53 peptide (3E peptide) and the control scrambled peptide. Upon doxycycline treatment the wound created in the control cells healed faster, suggesting that the R273H p53 mutant expressed indeed exhibited GOF properties, marked by higher proliferation and migration. The cells treated with the scrambled peptide showed similar closure time as the control cells, signifying that the scrambled peptide did not have any apparent pleiotropic effect. Interestingly, the cells treated with the 3E peptide phosphomimic p300-p53 interfering peptide migrated and proliferated slower than the control cells, implicating a possible role of p300 interaction in the tumorigenic potential of R273H p53 (Figure 6D). Moreover, the delay in wound closure observed upon the 3E peptide treatment was comparable to the untreated (doxycycline negative) cells, suggesting that the peptide treatment may have abrogated the effect of the p53 mutant expression. Furthermore, in the doxycycline negative cells the control, 3E peptide, and scrambled peptide-treated cells migrated and proliferated at similar rates suggesting that the effect observed upon 3E peptide treatment in the doxycycline-treated cells had a direct correlation with the presence of mutant p53 (Figure 6D). Collectively, these results implicate the direct role of p300 (and possible p300 autoacetylation) in the induction of R273H tumorigenic functions, creating a positive feedback loop between the two proteins. Western blotting analysis was done with the cells used in the assay to ascertain the expression status of R273H p53 in the presence and absence of doxycycline treatment (Supplementary Figure S7A). Since in the above experiment the mutant R273H was artificially expressed in H1299 cells, it was important to demonstrate a similar phenomenon in a cancerous cell line endogenously expressing the R273H p53 mutant. Therefore, a similar wound healing assay was performed in the AW13516 oral cancer cell line which expresses high levels of the R273H p53 mutant. Similar to the previous experiment, 3E peptide drastically impaired the rate of wound closure (Supplementary Figure S7B, middle panel, 12 hours) in comparison to the untreated and scrambled peptide treated controls (Supplementary Figure S7B, top and bottom panels, 12 hours).

**Figure 6:**
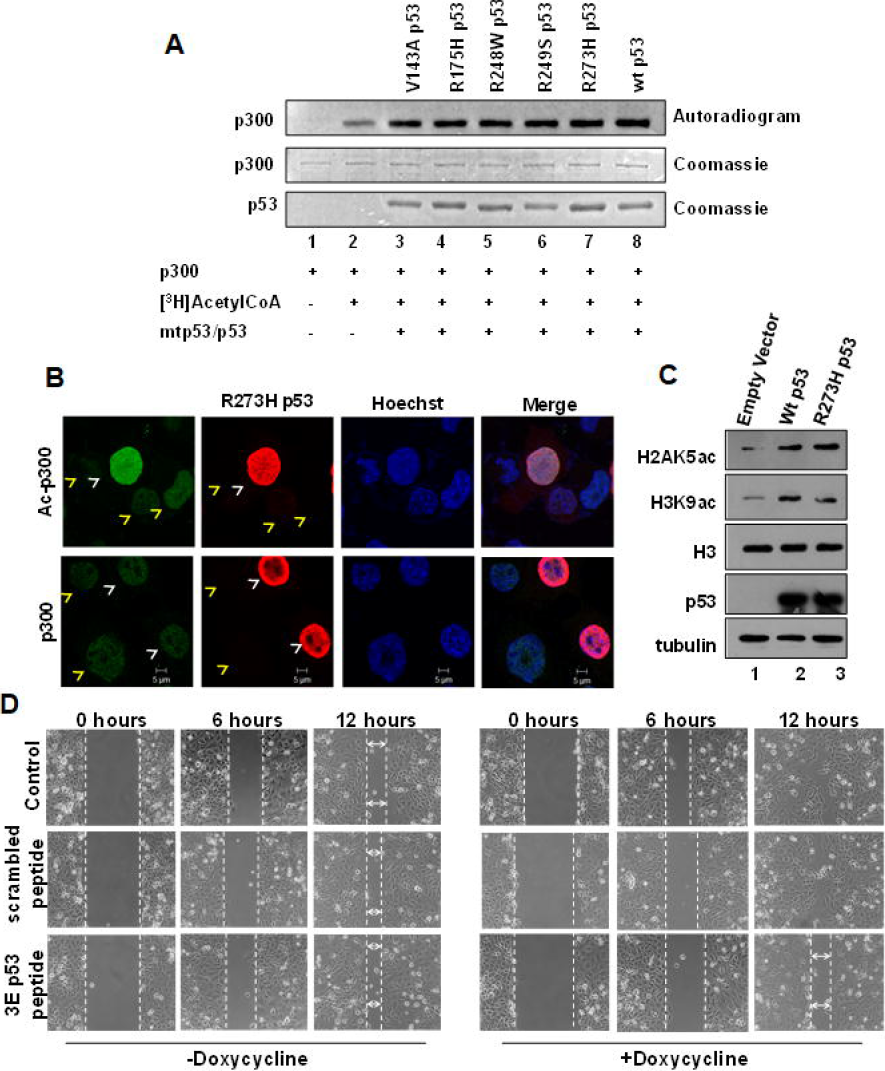
p53 Gain-of-Function (GOF) mutants are inducers of p300 autoacetylation. (A) An *in vitro* HAT assay was performed to determine the levels of autoacetylation of p300 in the presence of recombinant GOF p53 mutants and wildtype p53. (B) p53 null H1299 cells were transfected with R273H p53 mutant. The levels of Ac-p300 (autoacetylated p300) (top panel, green), p300 (bottom panel, green) and mutant p53 red were determined by immunofluorescence. Transfected cells and untransfected cells are indicated by white and yellow arrows respectively. (C) Western blotting analysis was performed to determine the levels of p300-specific histone acetylation levels in H1299 cell transiently transfected with empty vector, wild type p53, and the DNA contact mutant R273H p53. Histone H3 levels served as the loading control. (D) Representative images (of two biological repeats) of a wound healing assay performed in doxycycline-inducible R273H p53 H1299 stable cell line. The cells were treated with either phosphomimic p53 peptide or scrambled peptide as indicated with or without doxycycline treatment.

To further establish that the GOF mutant p53-mediated induction of p300 autoacetylation maybe an oncogenic pathway through which mutant p53 can exert its tumorigenic potential, we tested whether Loss-of-Function (LOF) tetramerization-defective mutants could also induce p300 autoacetylation. Tetramerization-defective mutants were transfected into H1299 cells and the levels of autoacetylated p300 was probed by immunofluorescence. It was found that the p53 tetramerization mutants failed to enhance p300 autoacetylation in the H1299 cells (Supplementary Figure S8A and S8B). It has been shown earlier that p53 proteins that cannot form tetramers, lose their ability to interact with p300 (64). It is fair to assume that the physical interaction of p53 and p300 is a key requisite for the induction of p300 autoacetylation. The L344A p53 tetramerization-defective mutant was tested in an autoacetylation assay. The autoacetylation of p300 does not alter in the presence of the mutant (Supplementary Figure S8C). When H1299 cells were transfected with the L344A p53 mutant, we observed no enhancement in the global levels of p300 autoacetylation or any alteration in p300 protein levels (Supplementary Figure S8D). These experiments revealed the importance of p53 tetramerization and the requirement for the direct interaction between p53 and p300 for p300 autoacetylation and augmentation of its acetyltransferase activity (Figure 7).

**Figure 7:**
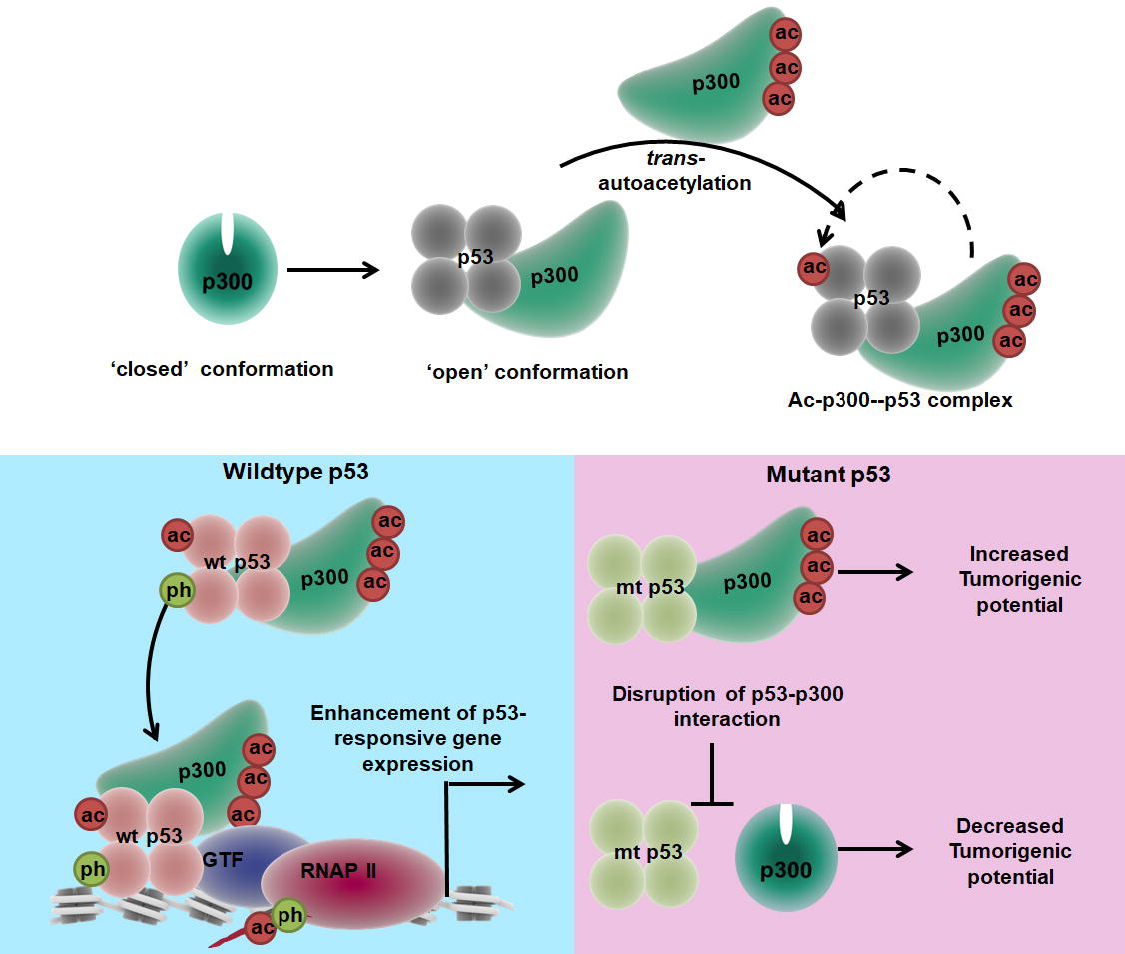
Model depicting the physiological significance of p53-mediated enhancement of p300 autoacetylation. The histone acetyltransferase is present in a ‘closed’ inactive conformation when present in an unbound state, but in the presence of p53, a substantial rearrangement in the domain architecture of p300 is observed. In the p53-bound complex, p300 undergoes a conformational switch from an inactive to an ‘open’ conformation. This open conformation state of p300 is more conducive to intermolecular autoacetylation, a phenomenon which enhances the catalytic activity of p300. When p300 is activated by wild type p53, there is a re-distribution of autoacetylated p300 onto p53-responsive gene promoters, leading to the downstream tumor suppressive functions of p53, whereas when mutant p53 induces the autoacetylation of p300, we observe an increased tumorigenic potential. The gain-of-function effect of mutant p53 is abrogated when the interaction between mutant p53 and p300 is disrupted.

### DISCUSSION

The functional crosstalk between p300 and p53 is well established; however, the role of p53 in the modulation of the enzymatic activity of p300 has not been addressed before. The ChIP-seq analysis of ac-p300 genome occupancy after p53 induction provided further insights into the transcriptional cascades triggered by effector (p53)-mediated induction of p300 autoacetylation. p300 autoacetylation can control its catalytic activity (12), but the dynamics of p300 autoacetylation under physiological stimuli are not fully understood. The present findings provide experimental evidence that establishes p53 as a *bona fide* modulator of p300 autoacetylation. Remarkably, among all of the substrates of p300 tested, only p53 enhanced p300 autoacetylation, thus confirming the molecular specificity of this phenomenon. The preferential induction of p300 autoacetylation by p53 over that of other acetyltransferases further exhibited the specificity of p53-mediated induction of p300 autoacetylation. This result further suggested that the induction of autoacetylation is not merely a by-product of enzyme-substrate interaction. No structural information currently available regarding the full-length p300 and tetrameric p53 complex. By using cryo-EM microscopy, we resolved the structure of p300 and p53 tetramer to gain insights into the molecular mechanisms of p53-induced allosteric activation of p300. Earlier structural studies have shown that the p300 core catalytic domain exists in a compact spatial conformation with basal catalytic activity. In the ‘un-activated’ conformation, the RING domain forms an obstruction to substrate-binding by occluding the substrate-binding groove of the HAT domain, which is buried inside the compact structure; importantly, this model was corroborated by our cryo-EM structure (59). In the p53-p300 complex, however, a distinct alteration of p300 conformation was observed. In this structure, the inhibitory RING domain was displaced away from the substrate-binding pocket of the p300 HAT domain in a conformation that could be characterized as ‘open’ and ‘activated’. These cryo-EM structural studies suggest that the ‘open’ conformation of p300 precedes the induction of autoacetylation, which is required for the maximal activation of p300 lysine acetyltransferase activity. The structure of the complex presented here indicated that N-terminal domains of p53 interact first with p300 to induce conformational changes and thereby the C-terminal domains of p53 gain access of the HAT domain of p300 (Supplementary Figure S4) suggesting the activation mechanism is allosteric in nature. Therefore, these data indicated that the ability of an effector to modulate the p300 conformational state can directly affects p300 acetyltransferase activity. p300 is recruited to gene promoters by several transcription factors (10, 65) where it can assemble transcription complexes that facilitate gene expression. The autoacetylation of p300 has been proposed to induce distinct structural changes that are critical for its activity. Black *et al*. have shown that the presence of p300 at the PIC exerts an inhibitory effect of transcription *in vitro*. The release of p300 from the complex is attributed to the possible conformational switch in the p300 structure after autoacetylation (66). It is clear from this study that p300 autoacetylation is essential for maximal transcription to proceed. The predominant occupancy of ac-p300 on the TSS of the promoters is a novel finding based on the obtained ChIP-seq analysis. These data are in agreement with the factor-dependent recruitment of the acetyltransferases p300/CBP and subsequent gene activation, as previously reported. As shown from our biochemical and structural data, the direct interaction of p53 with its coactivator p300/CBP is functionally important in this process and serves as the mechanism of induction of autoacetylation. Ceschin *et al*. have shown that CBP is methylated at multiple sites by the arginine methyltransferase protein CARM1 and the arginine methylation of CBP stimulates CBP activity through the induction of CBP autoacetylation. Use of polyclonal antibodies recognizing specific CBP methylation species, has indicated that each methylated class of CBP has differential HAT activities and distinct transcriptional effects, thereby diversifying the estrogen receptor response (67). The contextual recruitment of p300 has been noted in another study performed in the MCF7 cell line, in which it has been observed that in resting cells, p300 appears to show a preference towards neural lineage gene promoters, whereas the enrichment of p300 shifts toward estrogen-regulated gene targets after estradiol treatment (68). In a recent report, CBP has been shown to interact with RNAs. At active enhancers, CBP interacts with enhancer RNAs, which bind to the CBP catalytic domain, inducing CBP autoacetylation (69). The resultant enhancement in CBP activity leads to increased histone acetylation and expression of target genes. In the present work, the differential occupancy of p300 and ac-p300 was investigated after p53 activation. After autoacetylation, in the presence of p53, p300 appeared to show a strong preference for the TSS (Figure 5A and Supplementary Figure S5A), in agreement with the recruitment model discussed earlier. The binding of ac-p300 was strongly associated with RNA Pol II occupancy and the promoter acetylation mark H3K27ac. This finding suggested that ac-p300 is indeed involved in the assembly of the basal transcription machinery at promoter-proximal regions of genes. The overall pool of p300 did not show an appreciable alteration in chromatin occupancy in the presence of p53, whereas a distinct redistribution in ac-p300 enrichment was observed. Notably, the integration of the ChIP-seq data with the gene expression microarray analysis revealed that the enrichment of acp300 was a better determinant of p53-mediated gene expression than the presence of unmodified p300. This result was in agreement with previous studies in which p300 has been shown to occupy transcriptionally silent regions or the chromatin, whereas the co-occupancy of (presumably autoacetylated) p300 with its histone acetylation mark H3K27ac was observed at transcriptionally active genes (70). With the current evidence, it can be speculated that the ability to induce structural alterations in p300, leading to enhancement of its acetyltransferase activity, may be a key deciding factor in p300 chromatin recruitment and downstream transcriptional programs. The ChIP-seq data suggest that the phenomenon of factor-induced p300 autoacetylation may play a pivotal role in the integration of stimulus-driven transcriptional pathways. Specifically, the recruitment of the catalytically active form of p300 may trigger a rapid transcriptional response to internal and external signaling cues. In this study, it is the p53-driven pathway that stimulates a burst in ac-p300 levels and an alteration of the epigenetic landscape, which may be essential for these p53 signaling pathways. We hypothesize that after p53 is stabilized in the cell or overexpressed, it associates with its ubiquitously expressed co-activator p300. This association triggers an allosteric conformational switch in p300, thereby enhancing intermolecular autoacetylation. It is this catalytically active form of p300 that is subsequently enriched in p53-target gene regulatory chromatin regions via its interaction with p53. The active, ac-p300 then acetylates histone tails at these loci, decompacting chromatin and leading to enhanced transcription p53-dependent and (possibly) independent gene-networks (Figure 7, Table S2).

A large proportion of tumors harbor mutations in p53 that nullify specific DNA-binding properties. A subset of these mutants, the gain-of-function mutants, also gains aggressive proliferation properties, and it is now known that this is achieved by relocation to regulatory regions of genes responsible for proliferation and drug resistance followed by activation of expression of those genes. The results presented here underlines that in spite of the loss of DNA-binding properties, these mutants retain their ability to allosterically activate the coactivator p300, thus promoting activation of these growth promoting genes in the new locale. Abolition of the gain-of-function properties by a site-directed peptide inhibitor of p53-p300 interaction suggests that these classes of inhibitors may offer promise against tumors bearing aggressive gain-of-function mutant p53 alleles (Figure 7).

The activation of p300 by p53 suggests that p300 may be preferentially activated when it is recruited to p53 at its target sites. This allosteric activation would suggest an enhanced activity of p300 when it is complexed with target-site bound p53, thereby leading to localized transcriptional initiation. Thus, the augmented activity of p300 at genes to which it is recruited by p53 adds another element of specificity beyond that of the transcription factor binding. Moreover, p53 was able to modulate the levels of histone acetylation concomitant with the induction of p300 autoacetylation, thus signifying that the tumor suppressor p53 can alter the epigenetic landscape through the regulation of p300 acetyltransferase activity. We speculate that the allosteric activation of p300 by p53 demonstrated here may be a representative example of a general mechanism for attaining the high degree of spatiotemporal specificity required for attaining exquisite and intricate gene regulation.

## ACCESSION NUMBERS

The ChIP-seq data for p300 and ac-p300 is available at NCBI, SRA Accession ID: SRR5831017. The ChIP-seq data for RNA Pol II, H3K27ac, and H3K4me1 were retrieved from Data Bank of Japan (DDBJ), accession number: DRA001860 (http://dbtss.hgc.jp/) (71). Microarray of inducible p53 H1299 cell line data: GEO: GSE57841 (25). The cryo-EM density maps and coordinates of p300-p53 complex and free p300 were deposited in the Electron Microscopy Data Bank (EMDB) under the accession numbers EMD-6791 and EMD-6792, and deposited in the RCSB Protein Data Bank (PDB) under the accession codes 5XZC and 5XZS, respectively.

## ACKNOWLEDGEMENTS

We acknowledge Mr. Madavan Vasudevan and Ms. Madhura Tathode, Bionivid Technology Pvt. Ltd., for the ChIP-seq analysis. We thank Mr. Chiranjit Biswas for cryo-EM data collection and Mr. Sayan Bhakta for troubleshooting during image processing. We acknowledge Dr. Dirk Gründemann (University of Cologne, Germany) for the pEBTetD SLC22A1 construct. We have used illustration templates from the website somersault1824 (http://www.somersault1824.com) available under a Creative Commons Attribution-Noncommercial-Share Alike license (CC BY-NC-SA 4.0) in the Model (Figure 7) and Graphical Abstract.

